# PoseR - A deep learning toolbox for classifying animal behavior

**DOI:** 10.1101/2023.04.07.535991

**Authors:** Pierce N Mullen, Beatrice Bowlby, Holly C Armstrong, Angus Gray, Maarten F Zwart

## Abstract

The actions of animals provide a window into how their minds work. Recent advances in deep learning are providing powerful approaches to recognize patterns of animal movement from video recordings using markerless pose estimation models. There is an increasingly rich field of unsupervised and supervised methods for classifying animal behavior built upon the outputs of pose estimation models. However, these methods often rely on species and task-specific feature engineering of trajectories, kinematics and task programming. Highly generalized solutions that use only pose estimations and the inherent structure of animals and their environment provide an opportunity to develop foundational, contextual and, importantly, standardized animal behavior models for efficient and reproducible behavioral analysis. Here, we present PoseRecognition (PoseR), a behavioral classifier leveraging action recognition models using spatio-temporal graph convolutional networks. We show that it can be used to classify animal behavior quickly and accurately from pose estimations, using zebrafish larvae, *Drosophila melanogaster*, mice, and rats as model organisms. PoseR can be accessed using a Napari plugin, which facilitates efficient behavioral extraction for bout-like behaviour, annotation, model training and deployment. Our tool simplifies the behavioral analysis workflow by transforming coordinates of animal position and pose into semantic labels with speed and precision. Furthermore, we contribute a novel method for unsupervised clustering of behaviors and provide open-source access to our zebrafish datasets and models. The design of our tool ensures scalability and versatility for use across multiple species and contexts, improving the efficiency of behavioral analysis across fields.

## Introduction

Decoding animal behavior from video recordings allows us to understand its neural underpinnings. Identifying where an animal is, i.e., its position and pose, during video recordings has largely been solved by advances in deep learning, but recognizing animal movements or sequences of poses as meaningful behavior remains a more difficult problem for neuroscientists (1,2). Existing analysis workflows fall into two categories: unsupervised behavioral discovery or supervised behavioral classification. Unsupervised discovery leverages machine learning models to identify distinct actions of animals from either raw videos or pose estimations requiring no prior knowledge (3–9). This approach is useful for researchers looking to discover new behaviors but requires thorough post-hoc analysis and is sensitive to parameter selection and subjective interpretation of the number of clusters. As a result, it may lead to inter-lab discrepancies within the same dataset.. Alternatively, supervised classification applies prior knowledge of the behaviors that a researcher would like to extract from video recordings and teaches them to machine and deep learning models to improve the efficiency of analysis (10–16). The advantage of a generalized classifier is that there is no requirement for generating new embedding spaces of pose and features or aligning new data to pre-existing latent spaces; classifiers relying only on pose can be used in a plug-and-play fashion making them more accessible to the research community. Previous classifier approaches have typically focused on specific contexts of behavior of mice, fruit flies and, to a limited extent, zebrafish, often requiring the pre-computation of species-specific features or task programming (16,17). To produce a classifier architecture that doesn’t require extensive feature or task engineering and generalizes across contexts, backgrounds and species, we propose representing animal pose as a skeleton-like graph upon which temporal and spatial relationships between nodes can be learned to accurately predict behavior.

To this end of classifying diverse animal behavior, we utilized skeleton-based action recognition deep learning architectures that have demonstrated success in human action recognition(18). These architectures use graph neural networks to learn both the spatial and temporal components of pose estimations, treating them as nodes of a graph upon which convolution can be applied. Graph neural networks are revolutionizing our ability to model complex relationships between connected components with breakthroughs in solving the protein folding problem, recommendation systems, and drug discovery (19–21). A graph consists of nodes and edges where edges describe the relationship between nodes. Nodes can contain features, for instance (x, y) coordinates and confidence interval of a pose estimation. Mathematically, the pose graph is described as G = (V, E), where V represents the body part nodes and E the edges corresponding to the anatomical relationships between body parts. Convolutional operations on graphs involve aggregating feature information for each node from neighbor nodes to produce a new feature representation of the pose graph where knowledge of the state of nearby nodes contributes to the state of each node. For every time point in a behavior, the pose graph can be transformed in this way and the temporal features of these aggregated nodes can then be learned to classify behaviors.

### Design and Implementation

Our strategy to develop a new behavioral classifier based on these models was to simplify and accelerate three main steps in the behavioral analysis pipeline: 1) extraction, 2) annotation and finally 3) classification of behavior. We first applied signal processing methods to identify windows in which behavior is occurring, enabling rapid annotation of thousands of behaviors. The coordinates of the points on an animal body extracted using pose estimation, for example by DeepLabCut (1) or YOLO (22), are then used in space *and* time in spatial-temporal graph convolutional networks. We included easy-to-use functions and an accompanying tool, called PoseRecognition (PoseR), as an open-source plugin for the popular multi-dimensional data viewer Napari (23,24) to make training and deploying these deep learning models more accessible to a wider audience, and for the performance and visualization benefits it offers.

We used zebrafish larvae to first develop an efficient behavioral analysis workflow and then demonstrate its general use when applied to other species, multi-animal social contexts, and when including environmental context. Zebrafish exhibit a large repertoire of behaviors encompassing environmental exploration, escape, and predation (25). Click or tap here to enter text.Previous work to develop classifiers of zebrafish behavior has focused on binary classification of prey-capture swims versus non-prey capture swims in a fixed-head preparation (28,29). Several studies have developed unsupervised methods to uncover zebrafish swim types of freely swimming zebrafish; K-means clustering was used to identify 15 swim types from swim kinematics (30), density-based clustering was used to identify 13 swim types from kinematics (27), the FuzzyART algorithm was used to reveal around 50 clusters from trajectories of adult zebrafish (31), hierarchical clustering of four kinematic parameters was used to reveal 3 prey-response swims (32), autoencoder latent-embedding coupled with principal component analysis and dimensionality-reduction of spectrograms was used to obtain 22 clusters from raw video (33), tSNE dimensionality reduction of 220 swim bout features was used to obtain 36 swim types from videos recorded at 60 frames per second (26), and, finally, independent-component analysis was used to obtain 4 swim types (34). These approaches further highlight the variability in the number of swim types produced from unsupervised methods. Of these datasets, none met all the criteria of 1) being openly accessible, 2) containing raw pose estimates and more than two behavioral classes, and 3) having high framerates (>60 fps). W therefore sought to produce and open source our own large, high-framerate dataset of zebrafish larval poses and swim types using novel unsupervised methods to test the accuracy of graph convolutional classifiers in learning zebrafish behavior. It is possible to elicit a wide range of zebrafish behaviors in an experimental environment with visual projection of choice stimuli such as the looming shadow of a predator, the random walk of small prey, or ebb and flow in a natural scene of a riverbed (26,27). We acquired high-speed videos (330 fps) of freely swimming zebrafish larvae under these conditions and tested the effectiveness of our end-to-end behavioral analysis toolbox PoseR in two scenarios: generating and classifying a small manually curated dataset and classifying a large novel unsupervised-generated dataset of zebrafish behaviors. We also tested PoseR on a mouse open field dataset (14) Click or tap here to enter text.a pre-clustered zebrafish dataset containing tail angles (27) Click or tap here to enter text.and mouse, fly, and rat datasets of social behavior (35–37) to demonstrate the applicability of this approach to the analysis of other species and species-specific social and context-dependent behaviors.

### Behavioral setup

All procedures were carried out according to the UK Animals (Scientific Procedures) Act 1986 and approved by the UK Home Office. Zebrafish larvae (4-7 <u>d</u>ays <u>p</u>ost fertilization (dpf)) were placed in an acrylic recording chamber (25 x 25 x 25 mm) containing system water. Visual projections were displayed onto diffuse acrylic beneath the recording chamber using a cold mirror (Edmund Optics, 45° AOI, 50.0mm Square, Cold Mirror, #64 451) and a projector (Epson EF-11 3LCD, #0011131458). The zebrafish larvae were illuminated using a custom infra-red LED (850nm) array beneath the chamber and recorded at 330 frames per second using a Mikrotron camera (MC1362) and a high-speed frame grabber (National Instruments, PCIe-1433). Images were acquired and dynamically cropped in Bonsai (40) and zebrafish larval positions were extracted using background subtraction and thresholding. This allowed for closed-loop presentation of stimuli based on the position and orientation of the larvae using the BonZeb package (32) and reduced file size to permit continuous recording to disk of long-duration videos. In some experiments, live low-saline rotifers were added to the imaging chamber to record zebrafish larval swims in the presence of prey.

### Pose estimation

A ResNet50 neural network was trained using DeepLabCut (1) to estimate the position of 19 points on the zebrafish body. Each eye was represented by 4 points and the remaining 11 points were positioned at equal intervals along the zebrafish midline from nose to tail fin (see Fig. 1). The neural network was trained and videos were analyzed using a Tesla V100 Nvidia GPU at the Kennedy High Performance Computing Cluster, St Andrews.

**Figure 1.**
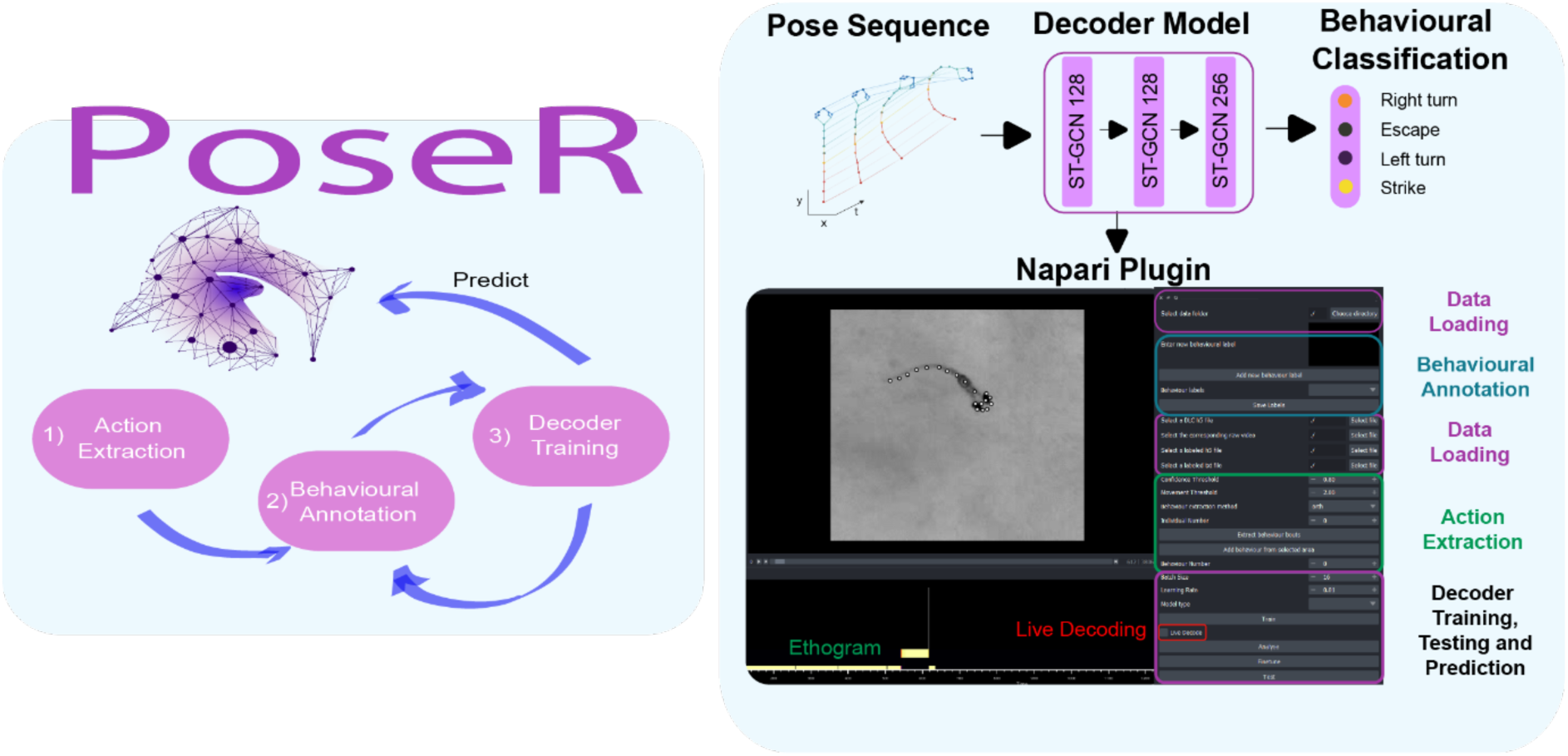
PoseR – A deep learning behavioral classification toolbox. A napari (24) plugin designed to accelerate extraction, annotation and training deep learning classifiers to predict animal behaviors. It combines popular deep learning packages Pytorch(38) and Pytorch Lightning(39) to simplify the process of training deep learning behavioral classifiers for a wide range of applications.

### Swim bout extraction

Due to the discrete bout-like nature of zebrafish swimming behavior it was relatively straightforward to define and extract periods in which behavior was occurring. Leveraging the lateral movement of the tail during swimming, the side-to-side motion of each body part was calculated, and a peak finding algorithm (41) was used to identify peak lateral movement and define the start and end of that swim bout. The side-to-side motion was calculated by first subtracting all coordinates by a center node (node 13) to get an egocentric representation of the zebrafish larval pose. The Euclidean trajectory and orthogonal trajectory for each node for each frame was calculated and future trajectories were projected onto the preceding orthogonal axis by dot product. This quantified the degree of perpendicular (side-to-side) motion of each node relative to the nodes’ previous position. The median side-to-side motion was smoothed with a gaussian filter (width = frames per seconds / 10). This representation of lateral motion was then thresholded using a median absolute deviation of 2 to extract peaks and windows around putative swim bouts using the Scipy find peaks function. Post-processing of these windows ensured swim bouts did not overlap and that the pose estimation confidence scores during that window were greater than 0.8.

### Manual labelling of swim bouts

All videos and DeepLabCut pose estimation were loaded into the PoseR napari plugin to facilitate easy swim bout extraction and manual labelling of behaviors. Swim bouts were extracted within the plugin as described above and video clips of those swim bouts could be cycled through in PoseR with pose estimations overlayed. Labels are assigned in the plugin from a drop-down menu and saved to an hdf5 file format to store the individual id, pose estimation, swim bout number, behavioral label and confidence scores. For initial validation, swim bouts were labelled as either left, right or forward swims, resulting in a manually classified dataset of 4368 swim bouts.

### Swim bout unsupervised clustering

34,015 swim bout pose estimations were extracted from behavioral recordings, resulting in an N x C x T x V array, where N is the number of swim bouts, C is the number of channels (X coordinate, Y coordinate and confidence interval of estimation), T is the number of timepoints in the swim bout, and V is the number of nodes on the zebrafish larvae body that were estimated. Swim bouts were aligned to the vertical axis ensuring all larvae were orientated facing north and modified to an egocentric coordinate system by subtracting the coordinates of a central node. The change in angle with respect to the central node during the swim bouts was computed for each node, resulting in an N x T x V array containing angle changes. The dimensionality of this array was reduced using tensor decomposition (42–44), where the data is approximated by a model consisting of a sum of components, with each component described by the outer product of three rank-1 tensors in the swim bout (N), time (T) and node (V) direction. This decomposition results in three matrices; a swim bout (N) x components factor matrix, a time (T) x components factor matrix and a node (V) x components matrix. The swim bout factor matrix contains a description of each swim bout according to the 10 tensor components. We used 10 components as this resulted in a low reconstruction error of 0.17, where the sum of components approximated the original dataset with an accuracy of 83%, whilst retaining stability in multiple replicates. Hierarchical agglomerative clustering (Scikit-Learn) was subsequently performed on this matrix where 30 distinct swim bout types resulted in a relatively low Davies-Bouldin score (45) and high silhouette score (46) in cluster evaluation.

### Spatial temporal graph convolutional network

A spatial temporal graph convolutional network (ST-GCN)(18,47) was modified into a Pytorch-Lightning module and the final architecture optimised using the Pytorch ecosystem across multiple Nvidia Tesla V100 GPUs (Kennedy HPC, St Andrews). The network was further modified to be shallower and wider consisting of three spatial temporal hidden layers of width 48, 256, 256, trained using a cross-entropy loss function and ADAM optimizer. All datasets were split into a training set (70%), validation set (15%) and testing set (15%) using sklearn where train, validation and test splits were not already provided in open-source datasets, and accuracy, precision, F1 score, and recall were calculated to evaluate the performance of each model.

### Installation

PoseR is installable via pypi https://pypi.org/project/PoseR-napari/ or via GitHub https://github.com/pnm4sfix/PoseR.

## Results

### PoseR enables fast coding and decoding of a small set of behaviors

We initially aimed to develop and validate our toolbox by rapidly coding a small subset of trivial zebrafish behaviors (left, right and forward swims) and designing an ST-GCN model capable of correctly decoding these. Whilst in practice classifying left vs right swims is easily solved using classic signal processing techniques, it provided an appropriate proof-of-concept to test our analysis pipeline and application of ST-GCNs to zebrafish behavior. Zebrafish larval poses, consisting of 19 coordinates on the larval head, trunk, and tail, were extracted from video recordings using a DeepLabCut ResNet50 model and ∼4,500 swim bouts were manually labelled in the plugin according to the direction of the swim bout by a trained observer. The graph in ST-GCN models is a spatial and temporal representation of the animal upon which graph convolution can be applied to represent complex interactions between nodes in a pose. To examine how the spatial graph representation of pose changes over time during the frames of a video recording, each node extends and connects to its corresponding node for each video frame through time (Fig. 2A). These abstract spatial and temporal representations of an animal’s pose can be learned using a spatial temporal graph convolutional network and the subsequently trained model can be used to classify behavior in an experimental setting (18). We trained an ST-GCN network consisting of three spatial-temporal graph convolution layers for 26 epochs using early stopping to prevent overfitting to training data (Fig. 2A). Testing the model on the validation dataset resulted in a high average accuracy of 90% across swim types (Fig. 2B). We calculated a value of 0.9 for precision, recall and F1-score of the model’s predictions versus ground truth. Precision quantifies the ratio of true positives to total positive predictions whereas recall quantifies the ratio of true positives to the total true positives and false negatives. The F1-score is the harmonic mean of precision (does the model detect all the true positives) and recall (does the model detect the positive cases and only the positive cases). Applying the model to unseen swim bouts and plotting the distribution of heading angle changes during the bouts revealed tight distributions in the appropriate direction for left and right swims (Fig. 2C). No trajectory information was included with our dataset, which relied only on egocentric coordinates; however, our model was able to accurately classify forwards swims and resulted in a heading angle change distribution centered on zero degrees. This distribution was symmetric but wider and bimodal, suggesting sub-groups of forward swims. This initial left-right ST-GCN model provided a promising initial validation for our toolbox in predicting a small subset of manually labelled behaviors. We took this approach further to test the limits of the toolbox and produce a model capable of accurately classifying a wider, more diverse, and challenging range of zebrafish behaviors.

**Figure 2.**
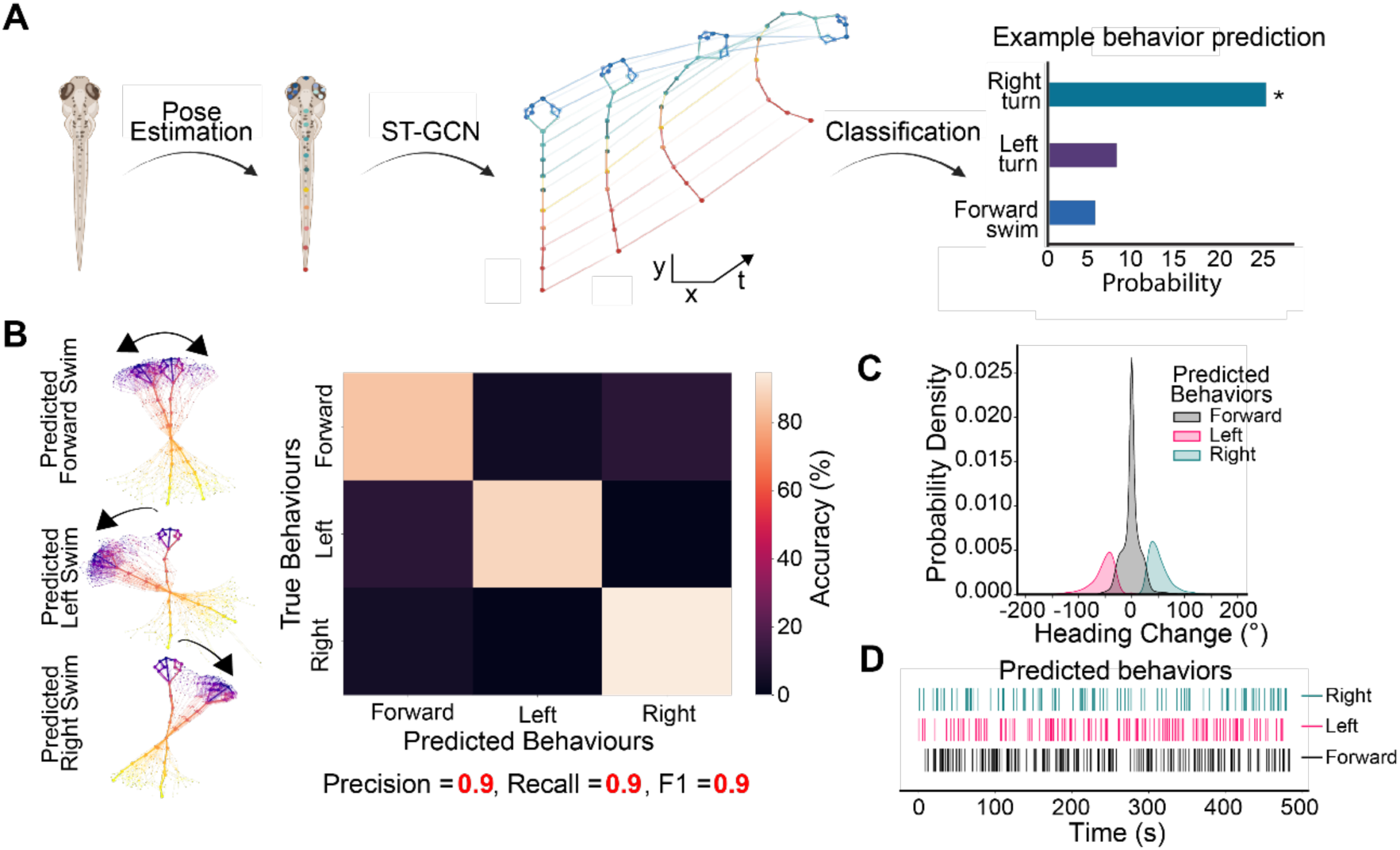
Deep Learning Behavior Classification Strategy. **A)** Graphical abstract created in BioRender (DJ27IPKD7A) demonstrating the workflow of classifying sequences of poses as zebrafish behavior. Zebrafish larval poses were extracted from videos using DeepLabCut. Sequences of poses can be represented as graphs with edges connecting body parts in space and time. **B)** These sequences were used to train a spatial-temporal graph convolutional neural network (ST-GCN) to classify swim bouts manually labelled as either left, right or forwards swims. Examples of correctly classified swim bouts are shown. **C)** Heading changes of unseen swim bouts classified by the trained left-right ST-GCN model. **D)** An example ethogram showing classified left, right, forward swims during a recording session.

### Generating a comprehensive zebrafish behavior dataset using tensor decomposition and agglomerative clustering

We next generated a larger dataset of zebrafish larval behaviors. We recorded long duration (40 minutes), high frame rate (330 fps) videos of zebrafish larvae behaving and responding to a wide range of closed-loop visual stimuli (32). These stimuli were chosen to elicit natural behavior such as escape reflexes, phototaxis and optomotor responses, resulting in a dataset of ∼30,000 swim bouts at high temporal resolution. We took a novel approach to clustering these swim bouts by first using tensor decomposition (42,43) to reduce the complexity of the dataset to 10 tensor components with each component containing a swim bout factor, body part factor and time factor (Fig. 3A). This resulted in a matrix describing the contribution of each swim bout according to the 10 tensor components, to which hierarchical agglomerative clustering could then be applied (Fig. 3B). To create a challenging dataset, we defined optimal clustering criteria as having a minimum cluster number of 15 and well-separated clusters with a silhouette score of > 0.2. This resulted in 30 distinct behaviors, which contained swim bouts that were homogeneous within each cluster and showed similarity to swim bout types that have been previously described (27,48) (Fig. S1). Symmetrical tail beats in clusters 2, 14 and 23 represented forward swims with increasing power from slow scoots to forward bursts. Broadly, clusters appeared to be initially separable by changes in heading direction with broad categories assigned left, right, forward, and large angle turns, which we mapped to a low dimension behavioral projection of each swim using uniform manifold approximation and projection (49)(Fig. 3C, Fig. S1A). Within these classes, the temporal dynamics varied depending on the vigor, amplitude, and number of tail oscillations and whether changes in direction occurred early or late within the swim bout. Clusters 9, 11, 15, 18, 22 and 27 appeared to show similarity with routine-turns described in the literature involving a change in orientation of about 40 degrees with no scoot (48). Burst-like swims were identified in clusters 1, 4, 6, 7, 12, 13, 23 and 28, where sustained large-amplitude cyclical tail oscillations were observed. Putative O-bend swims could be mapped to clusters 20 and 30 where an almost complete inversion of heading direction occurred with little evidence of large tail beats after the heading change. J-turn-like orientating swims showed similarity to cluster 3, 5 and 29 where the heading change is accompanied by small amplitude tail beats. Large amplitude escape-like swims were identifiable in clusters 16 and 24, where fish rapidly changed and swam with vigor in almost the opposite direction resembling slow-latency C-bend swim (SLC)(48). Long-latency C-bends (LLC), where the heading change and swim vigor were less extreme than SLC swims were seen in clusters 8, 10, 17, 19. Quantifying the presence of each swim type by visual stimuli highlighted the expected preference for left turns and right turns during the presentation of R-L and L-R optomotor gratings, respectively, as larvae aligned themselves with the direction of visual flow. Increases in swim activity across types were seen during the presentation of visual prey, with increases in forward swims when the prey was presented ahead of the larvae. Large amplitude swims were most prevalent in phototaxis, optomotor and prey presentation. In the presence of live rotifers, swim types 1, 3, 4, 5, 12, 14, and 23 dominated, representing a combination of burst swims and J-turns prevalent in zebrafish predation (Fig. 3D) (27,50–52). Using this novel approach, we succeeded in extracting and generating a large, complex dataset of a range of swim bout behaviors that zebrafish larvae employ during different visual contexts.

**Figure 3.**
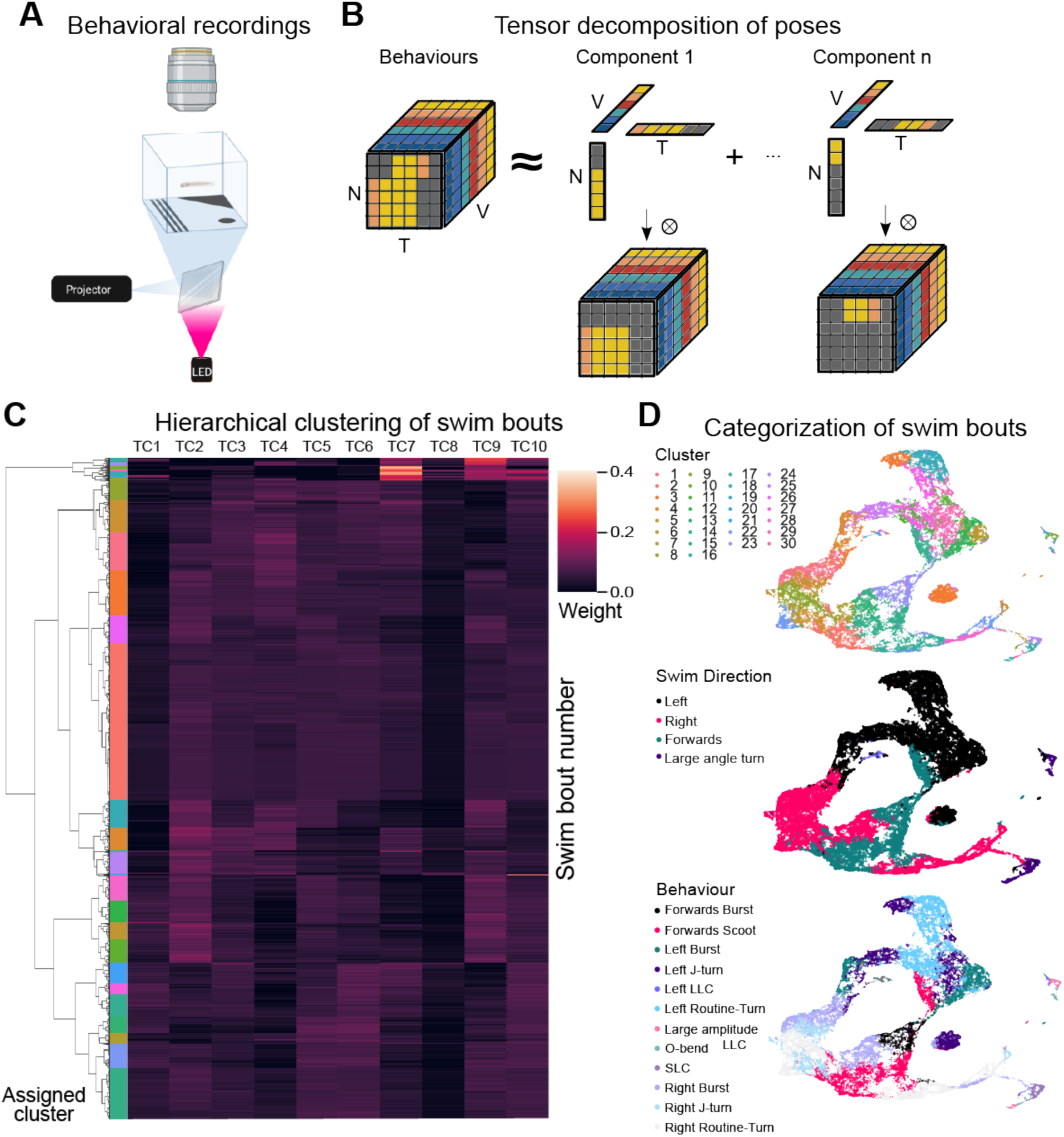
Clustering decomposed swim bouts. A) Graphic of the behavioral experimental setup for high-speed video recording and visual presentation, made in BioRender (II27IPKPNA) **B)** Tensor decomposition (PARAFAC/CANDECOMP) was applied to a 3d (rank-3 tensor) dataset containing egocentric node positions (in radians) of swim bouts, where the N dimension represented the swim bout number, the T dimension represented time and the V dimension represented the node/body part of the zebrafish. **C)** Agglomerative hierarchical clustering heatmap of the N x TC factor matrix showing how swim bouts could be grouped into similar clusters. **D)** A uniform manifold approximation and projection of the N x TC factor matrix color-coded by swim type cluster and broad swim class (left, right, forward and large angles swim).

### PoseR can be used to rapidly classify many complex behaviors

We next used PoseR to train an ST-GCN to recognize these more complex clustered swim behaviors with the aim of developing a universal zebrafish classifier that could be applied across a range of experiments. We found a high top-1 and top-3 unweighted-average accuracy (76% and 97%, respectively) for correctly classifying all behaviors in the test dataset. and this was achieved on a first run using PoseR’s built-in helper functions to optimize initial model hyperparameters. Top-1 and top-3 accuracy is often used to report the presence of correct behavioral classification in the top guess of the model (top-1) and in the top 3 (top-3) guesses. Model accuracy increased and loss decreased quickly during training with a batch size of 16, cross entropy loss function and ADAM optimizer before stopping early when the loss plateaued (Fig. 4A). The accuracy of the model’s initial predictions was evident by the bright band along the diagonal axis of the confusion matrix and a precision, recall and f1-score of 0.77, 0.76, 0.76, respectively (Fig. 4A, B). In addition to fast optimized training, we endeavored to evaluate the speed at predicting and analyzing new behaviors on different systems accelerated by either a GPU or a CPU. As expected, we found faster inference speeds on GPU based systems compared to CPU; with classifying of 1000 swim bouts taking approximately 20 seconds with a batch size of 10 on GPU systems. The latency to analyze one bout on an Nvidia Titan RTX GPU system was 2.85 ± 0.22 ms (Fig. 4B). The different frequencies with which swim types occur resulted in an imbalanced dataset where some swim types, in particular the large amplitude turns, were rarer. To address this issue, we included a weight calculation function in PoseR during training to estimate the best weights for each swim type derived from the rate of occurrence of that swim type relative to the total dataset size. This produced a model with a slightly higher accuracy of 77%, a more balanced precision, recall and f1-score for all swim types including less frequent swim types demonstrating the ability of PoseR to develop models to accurately predict behaviors in unbalanced datasets (Fig. 4B-D). Graphs are flexible data structures and graph nodes can be assigned multiple types of data, from (x, y) pose estimations to precalculated joint angles, or local video features. To demonstrate this flexibility, we trained an ST-GCN model on a large pre-clustered zebrafish dataset that contained only angle information of a zebrafish larval tail with no (x, y) pose estimations (27). This dataset contained 13 swim types recorded at 700 fps, and angle information for 8 nodes along the tail and tail nodes in the ST-GCN model were connected in a chain to represent the tail. Using PoseR, the trained ST-GCN model achieved top 1, 2, 3 accuracies of 88%, 98%, and 99% with an overall precision, recall and f1-score of 0.9, 0.88, 0.89, respectively (Fig. 4E), demonstrating the powerful application of this approach to classifying behavior from different types of data from a variety of sources.

**Figure 4.**
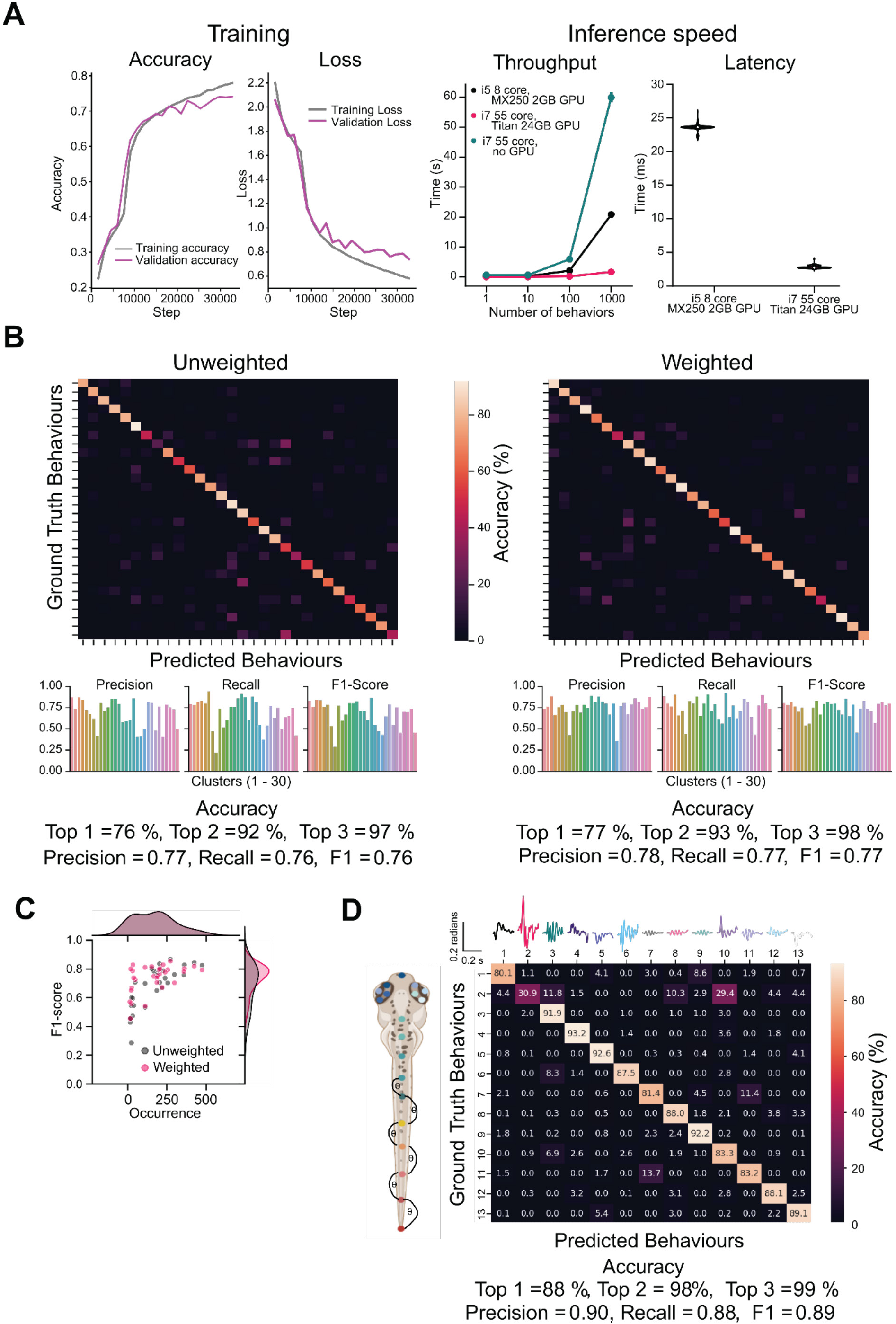
Accurate classification of zebrafish behavior using a trained ST-GCN model. **A)** Training and validation accuracy and loss reported during each training step. Latency for one sample and inference speed for batch sizes 1, 10, 100, 1000 on different systems: a CPU-only system, a system with 2GB MX250 GPU, and one with a 24GB Titan RTX GPU. **B)** Top: A confusion matrix highlighting the percent of correct predictions compared to ground truth behavioral labels for unweighted and weighted behavioral training data. Bottom: Precision, recall and F1-score for unweighted and weighted trained models. **C)** A scatterplot of F1-score versus the occurrence of a behavioral label to show weighting labels improves the overall F1-score. D) An ST-GCN model and subsequent confusion matrix trained on the ZebRep dataset (27,53) where only angle information is included in the feature matrix of the pose graph. The zebrafish graphic was created in BioRender (DJ27IPKD7A).

### PoseR can classify behaviors of other species and understand environmental context

PoseR can be extended to understand the patterns of movements of other animals, and within their environmental context. To demonstrate the utility of PoseR and the skeleton approach for training accurate behavioral models in this context, two ST-GCN networks were trained to classify three mouse behaviors recorded in an open field test (14), with one network trained in PoseR using pose information from the mouse body, and the other trained in PoseR using pose information from the mouse body and an additional five points demarcating the corners and center of the arena (Fig. 5A). This pre-existing dataset already contained manually labelled rearing and grooming behaviors, with the rearing behaviors sub-divided into supported, where the mouse leans on the arena to rear, and unsupported, where it does not (Fig. 5B). We excluded very rare ‘jumping’ labels for direct comparison with other tools that had removed these too. Behavior bouts were extracted from the dataset according to the labeling metadata and split into train, validation and test dataset splits using PoseR. Neural networks trained on body-only pose information excelled at identifying groom behaviors (92% accuracy) from rearing behaviors (100% accuracy), however due to the lack of environmental context within the pose estimation unsupported rearing was more often confused with supported rearing (Fig. 5C). Including information about the arena as nodes within the pose graph led to enhanced performance in recognizing and distinguishing supported and unsupported rearing and grooming producing a more balanced behavioral model with accuracies of 89%, 75%, 86% for support and unsupported rearing and grooming, respectively. Distinguishing between unsupported and supported rearing could simply be established post-hoc by the occurrence of the rear at the proximity of the wall, however here we use this example to demonstrate the versatility of ST-GCN models to include information about the environment to intrinsically make accurate predictions about behaviors distinguishable by the proximity of the occurrence to specific objects or features in the environment (Fig. 5D).

**Figure 5.**
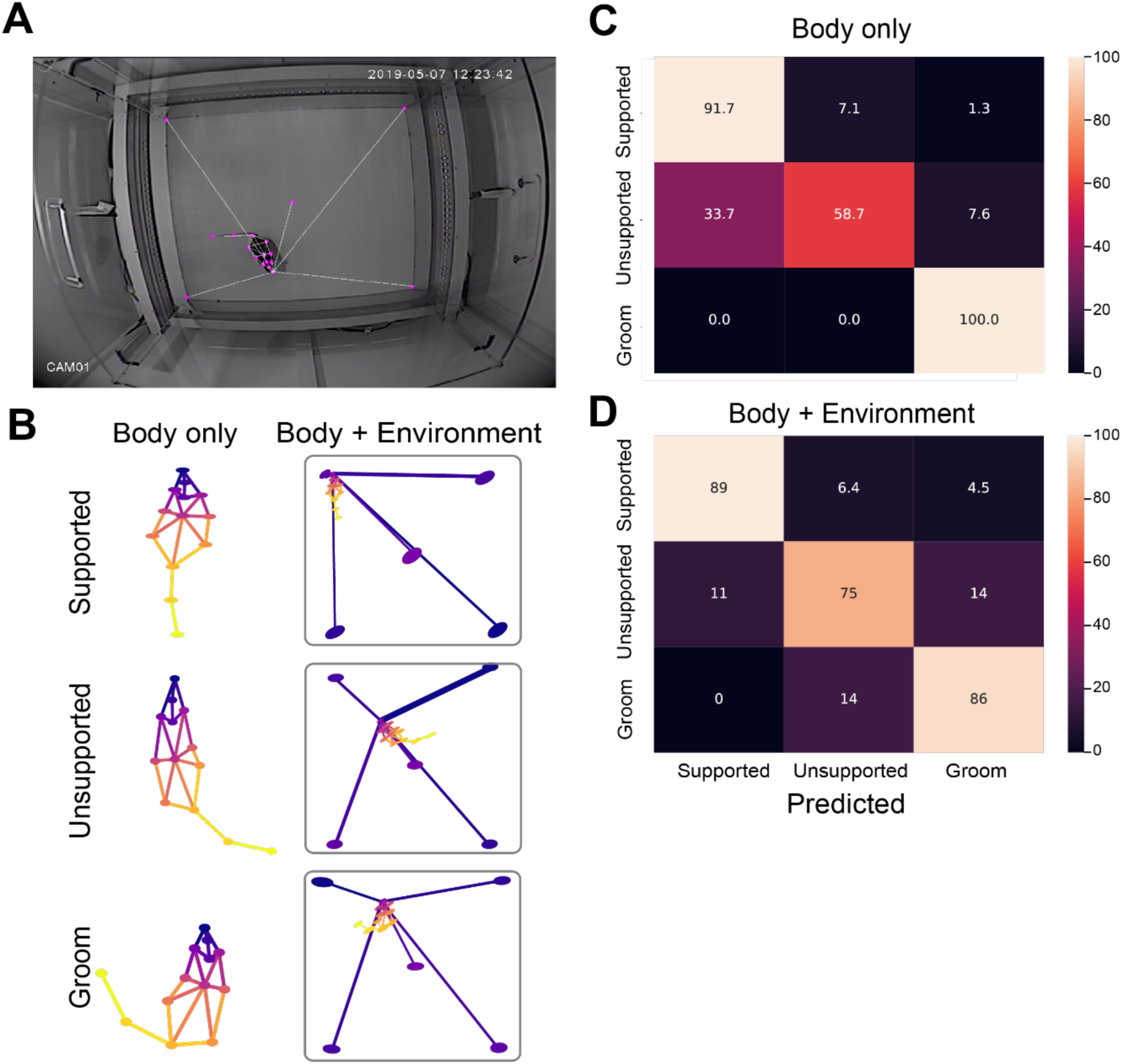
Accurate classification of mouse behavior in the open field test. **A)** The open field arena with tracked key points of the mouse body as well as the arena overlayed in magenta reproduced from Sturman et al under CC By 4.0. **B)** Dataset contains three behaviors: supported and unsupported rearing, and grooming; these were modelled with a body only or body plus environment spatial temporal pose graph. **C, D)** Confusion matrices showing the performance of an ST-GCN network trained on body-only pose information and body plus environment pose information, respectively.

### PoseR models can be extended to classify social behaviors

Engagement in social behaviors is an important metric for screening the effect of different pharmacological compounds and an efficient method of classifying these behaviors is crucial. In addition to including environmental context within the pose graph, multiple individuals can be represented and combined as individuals and/or fully-connected community pose graphs to learn social behaviors across different animal species. We used three published datasets, FlyvFly (35), CALMS21 (37), PAIRR24M (54)to demonstrate the versatility of PoseR and the ST-GCN approach in classifying social behaviors. Briefly, FlyvFly contains pose estimations of three points, the tip of each wing and the body center, of two flies interacting in a variety of contexts with behaviors labelled as lunge, wing-threat, tussle, wing-extension, circle, or copulation (Fig. 6A). PAIRR24M consists of twenty-five three-dimensional (3D) body coordinates of two rats interacting in an open field arena, which we used to demonstrate the ability of PoseR to train models of 3D pose sequences representing rat social behavior (Fig. 6B). CALMS21 is a dataset focused on four behaviors of two interacting mice, divided into two tasks defined by the number of annotators and size of training data available (Fig. 6C). We extracted pose sequences centered on each frame, with a window size of 14 frames. Pose estimations were assigned into train, validation and test splits and PoseR was used to train ST-GCN models to classify social behaviors. Models were able to accurately classify behaviors with respective accuracies of 98%, 85%, and 62% in the test splits of FlyVFly, CALMS21 and PAIRR24M, respectively, with example ethograms showing a favorable comparison between frame-by-frame ground truth and model prediction. The ethograms highlight a good precision of behavior across species and contexts, that is, the model accurately predicts when a behavior does occur. However, there were several instances of false positives, where the model predicts a brief occurrence of behavior when it does not occur, thus, some post-hoc and annotator led refinements of these short false positives are necessary for best results; these tools are included in the PoseR plugin (Fig. 6D).

**Figure 6.**
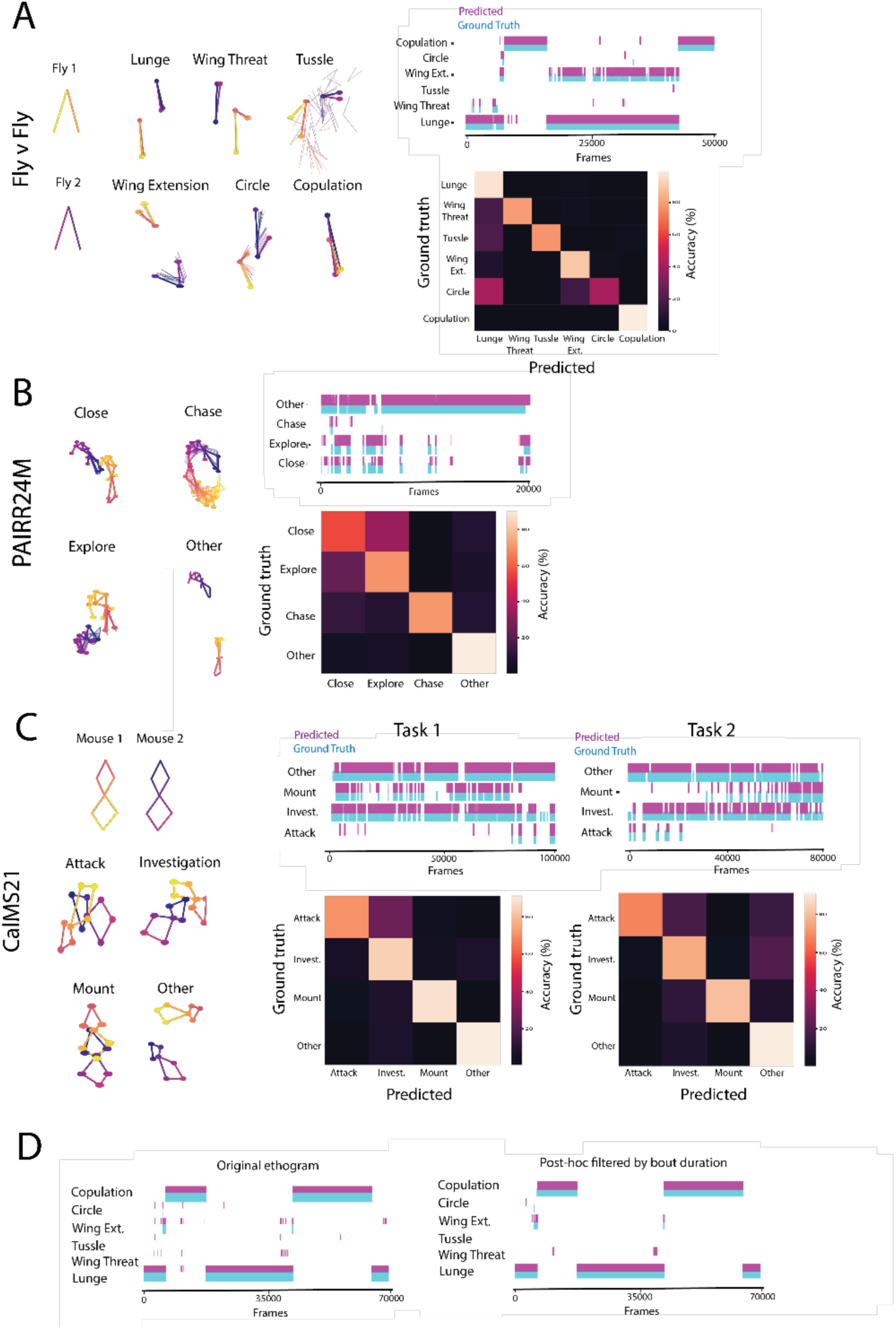
Classifying fly, mouse and rat social behaviors with PoseR. **A)** Fly v Fly dataset. Left: Each fly is represented by three nodes that label the tip of each wing and the center of fly. Middle: Six behaviors were labelled in this dataset; lunge, wing-threat, tussle, wing-extension, circle and copulation and appear distinct based on their temporal and spatial pose graph. Top right: An ethogram showing a frame-by-frame comparison of the ground truth behavioral label and model prediction. Bottom right: A confusion matrix denoting the accuracy of the model for each behavior. **B)** PAIRR24M dataset. Left: Each rat is represented by twenty-five nodes in 3D space. Social behaviors consisted of close, chase, explore, and other. Top right: An ethogram comparing the frame-by-frame accuracy of the model predictions versus ground truth labels. Bottom right: A confusion matrix highlighting the accuracy of the model for each behavior. **C)** CALMS21 dataset. Left: Each mouse is represented by seven nodes that label the body of the mouse. Four behaviours, attack, investigation, mount and other are contained in this dataset. Middle: Performance on task 1 visualised by ethogram comparing model predictions versus ground truth and a confusion matrix showing behavior level accuracy. Left: Task 2 containing reduced amount of training data, showing an ethogram example and confusion matrix. **D)** Left: Original ethogram of FlyvFly video subset, right: the same ethogram after post-hoc filtering defining the minimum bout duration for behavior to remove erroneous noisy predictions.

### Availability and Future Directions

Here, we presented a deep learning toolbox, PoseR, with the aim to accelerate the understanding of animal behavior. Using the versatile zebrafish larva as an animal model, we demonstrated the flexibility of PoseR in extracting and annotating behaviors from pose estimations and training deep learning models to recognize a range of these behaviors. We also highlight the versatility of applying tensor decomposition to behavioral data as a powerful method for dimensionality reduction preceding unsupervised behavior discovery. We designed PoseR to be fast and accessible: we leveraged Python libraries that offer user-friendly solutions to training deep learning models; use shallow networks that learn classification boundaries without adding computational overhead; and combined these tools into an established and responsive data viewer, napari. Our approach is validated by the rapid inference speeds and accurate models across species and behavioral contexts (Table 1). Models can be further refined within PoseR using finetuning functions to freeze and unfreeze select layers during training. This offers the ability to modify our pretrained models for animal behavior to new behavioral classes by, for example, training only the last classification layer of the model. We have demonstrated that PoseR is also inherently agnostic to animal species, relying only on pose estimations. Furthermore, researchers studying the interactions of animals with their environment and as a social group can include key point coordinates representing important features in the environment as well as connections between multiple individuals within our framework to understand how animals behave in a context-specific and social manner.

**Table 1.**
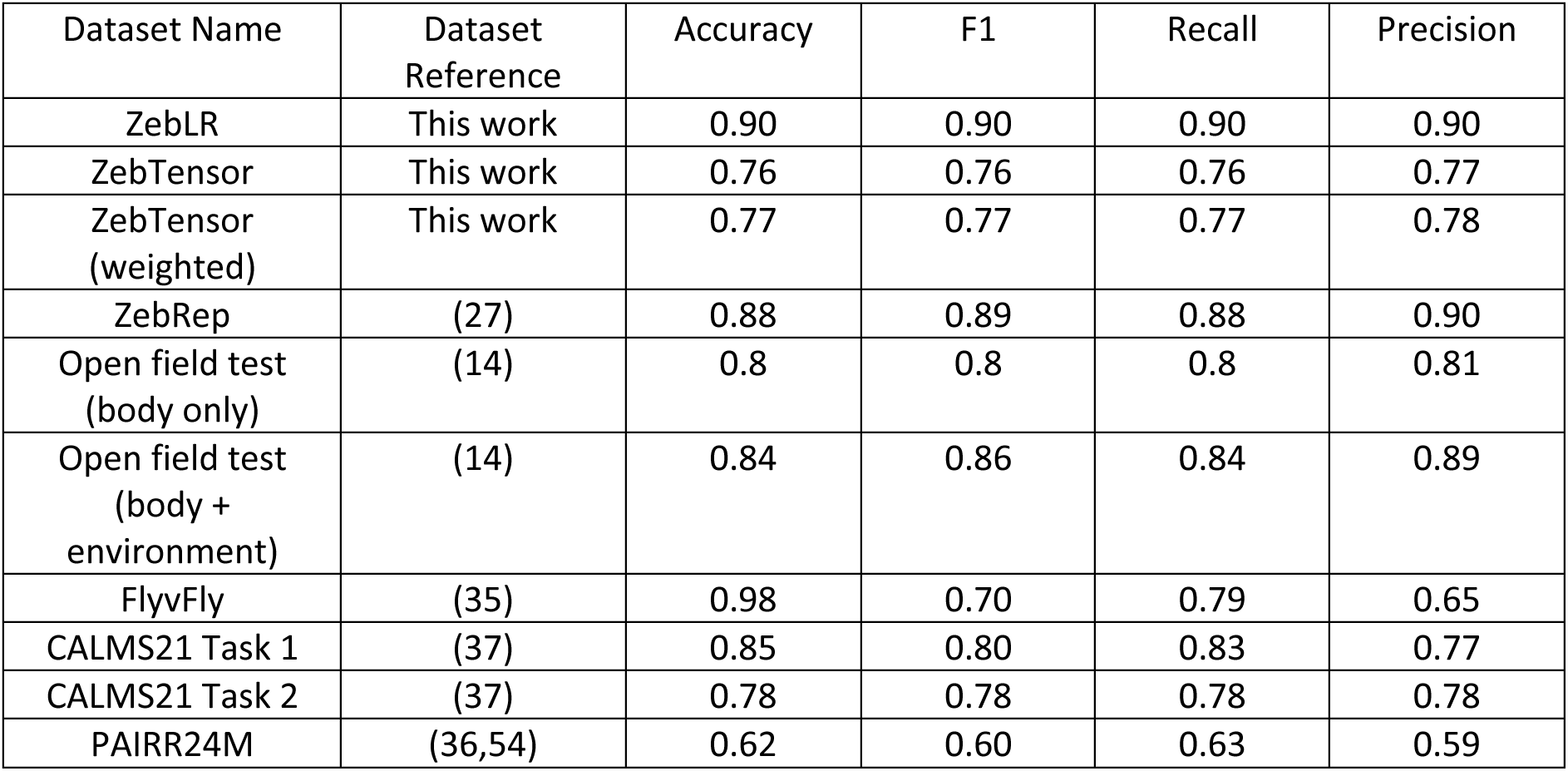
Accuracy metrics on datasets used to train models in PoseR. “Dataset” denotes the name used of the datasets used in this work, where “ZebLR” is the simple proof of principle left/right/forward swim dataset, “ZebTensor” is the tensor-decomposed clustered dataset of swim types to test the PoseR models on a large amount of behaviors, “ZebRep” refers to the published Marques et al. dataset, “Open field test (body only)” and “Open field test (body + environment)” refer to the Sturman et al. dataset with and without arena context. FlyvFly, CALMS21 and PAIRR24 are multi-individual datasets for social behavior. Accuracy, F1, Recall and Precision are averages across all behaviors within the specified dataset. In datasets where ‘Other’ is present as a label, this is excluded from the final metric score.

Whilst PoseR can be used with CPU-only systems, optimal throughput is achieved with GPU-based systems, drastically cutting inference speed and the time required to train a model. The current framework is built upon pose estimation outputs from the popular DeepLabCut Python package and we additionally provide pre-trained YOLO pose estimation models to facilitate plug-in-and-play pose estimation and behavior classification for the example species and contexts (22) (Fig. S3A). The classification models presented here are modified spatial-temporal graph convolutional networks. These networks have clear advantages: they require only pose information and confidence intervals of those estimations, and can simultaneously study the spatial and temporal components of these poses. Application of these networks to human action recognition datasets greatly improved classification accuracy in these challenges, and subsequently these types of models are beginning to be adopted in the study of animal behavior (55). (56)By including an individual and its environment in the open field test pose graph, we created a basic scene graph, where objects, individuals and their environment can be modelled as connected components. We further extended this to environments with multiple individuals; however, other key locations and objects could be included in a scene graph to provide more utility for researchers performing complex behavioral experiments involving interaction with other individuals and the environment.

Applying tensor decomposition provided a powerful way to extract distinct swim types and create a large behavioral dataset to train and test our supervised ST-GCN models. Previous attempts to extract zebrafish larvae behavior have used a range of techniques from t-SNE embedding, density-based clustering to FuzzyArt algorithms to effectively discover and describe zebrafish behaviors in a range of experimental conditions (26,27,31). However, unsupervised clustering, whilst a powerful and useful technique, does not represent an absolute truth of the number or separation of behaviors and can lead to an unreproducible and variable numbers of clusters depending on the context and parameter selection. We report a division of our swim bouts derived from visual stimuli and predation assays into 30 clusters. In similar experimental setups, others have reported a range of distinct swim types from 13 (27) to 36 (26). We created an ST-GCN model to accurately classify the 13 behaviors from (27) using only the angle information of the tail as no (x, y) pose estimation data was present in this dataset. The ability of the ST-GCN model to learn these different behaviors from one feature alone was testament to the versatility of graph neural networks and the well-clustered swim types in this dataset. However, our main aim was to teach ST-GCN models to recognize behavior from pose estimations, and we were unable to train an adequate model to learn the 36 behaviors from Johnson, *et al.*. We do not know whether this was due to the larger number of clusters, or as a consequence of the low temporal resolution (60 fps) of this dataset compared to our own (330 fps) and Marques et al. (700 fps). This provided the rationale for creating our own dataset, presented and openly accessible here, with high temporal resolution pose estimations of larval behavior evoked by a variety of visual stimuli and prey.

The PoseR toolbox developed here sits within a healthy ecosystem of emerging and maturing behavioral analysis tools that aim to discover behaviors in an unsupervised way or classify behaviors manually. PoseR initially aimed to provide a platform to efficiently extract zebrafish swim bouts and to train accurate behavioral classifiers to learn the repertoire of zebrafish behavior. However, we also found that using skeleton-based action recognition in PoseR approached the performance of classic convolutional models such as DeepEthogram in the context of the Sturman mouse open-field test (Table 3). Other tools such as BSOID (57) that specialize in open-field behavior classification achieve accuracies of around 90% on unsupported and supported rearing in bottom-up camera recordings in comparison with 89% and 75% presented here on the top-down Sturman dataset. Recall of groom behavior was significantly better when using pose estimations of the mouse body alone and performance was substandard when environmental features of the arena were included in the pose graph. It is therefore clear that balancing the inclusion of environmental context without hindering accuracy of non-environmental behavioral is crucial and further refinement in PoseR model architecture and training strategy is required to reach the high standard in this context (15,57). Notably, we observed comparable or slightly improved performance compared to TREBA (17) when PoseR models were trained on social fly and mouse datasets suggesting that PoseR may be more suited to these complex behaviors and contexts where the connectivity of the pose graph is important in the classification of the behavior (Table 2). An advantage of skeleton-based action recognition over video-based action recognition is that it is likely to be less susceptible to background noise in videos, assuming robust and mature pose estimation models are used to extract poses. Whilst pose estimation models and image-based action recognition models both suffer from background noise, skeleton-based action recognition models, as used here, can mitigate the effect of pose estimation errors by including confidence intervals of the pose estimation in the node features to teach the model when coordinates are confident or not. In our specific case, we acquired high frame rate (>300 fps) videos required for capturing zebrafish larvae behavior by dynamically cropping the region of interest around the zebrafish during acquisition to reduce image size and to save to disk in real time. This led to a dynamically shifting non-uniform background flow that could interfere with video flow-based methods of behavioral analysis. Where a camera’s field-of-view is not stable, whether during recording in the wild or, as in our case, where a camera’s field of view is cropped to track a fast-moving animal, we think the skeleton-based approach is likely to be more optimal than flow-based methods. The field of skeleton-based action recognition is one strand of the action recognition field as a whole and is making rapid progress with the development of novel and more powerful architectures for classifying behavior (51–53). It will therefore be interesting to see the comparative performance of these approaches on the contexts presented here. We observed that ST-GCN models for classifying behaviors required relatively short training durations, and propose this is likely due to the previously observed phenomenon of quicker convergence in graph networks(61). Across the datasets presented here, training duration ranged from 13 minutes to 3 hours; other tools like TREBA and DeepEthogram have reported 24 hours of training. By ensuring models are shallow we can also achieve fast inference speeds across datasets of between 240 and 435 fps, leading to analysis durations of under 2 minutes for hour-long recordings at 30 fps (Table 2). For zebrafish-specific data, analyzing extracted bouts only instead of individual frames reduces analysis durations, for example, for a 40-minute video recorded at 330 fps from 33 minutes to 9 seconds (Table 4). An important aim of neuroscience is to understand how neural activity underlies behavior and these inference speeds would be sufficient to enable real-time classification of behavior. Our tool, combined with fast pose estimation (62) and neural activity recording, therefore offers an exciting opportunity to directly correlate *in vivo* neural activity with behavior, and trigger experimental manipulation with behavioral cues, advancing the effort to understand how the brain produces behavior.

**Table 2.** Example durations for analysing poses from one video for each dataset model. For each dataset, an example video was analysed to demonstrate typical durations of analysis with a batch size of 16.

| Model | Video duration | # frames | Time taken |
| --- | --- | --- | --- |
| ZebLR | 40 minutes @ 330 fps | 780065 | 33 minutes |
| ZebTensor | 40 minutes @ 330 fps | 780065 | 32.6 minutes |
| Open field test | 10 minutes @ 25 fps | 15253 | 36.8 seconds |
| CALMS21 | 2 minutes @ 30 fps | 3632 | 3.5 seconds |
| FlyVFLy | 28 minutes @ 30 fps | 49406 | 18.3 seconds |
| PAIRR24M | 60 minutes @ 30 fps | 108000 | 1 minute 52.3 seconds |

**Table 3.** Comparison of PoseR against other behavioral tools. Macro-weighted F1 scores for PoseR models in comparison with reported results from DeepEthogram and TREBA on the Open Field Test, FlyvFly and CALMS21 datasets.

| Dataset | PoseR | DeepEthogram (medium) | TREBA |
| --- | --- | --- | --- |
| Open field test | 0.84 | **0.92** | - |
| FlyVFLy | 0.70 | - | **0.72** |
| CALMS21 Task 1 | **0.80** | - | **0.83** |
| CALMS21 Task 2 | **0.78** | - | 0.77 |

**Table 4.**
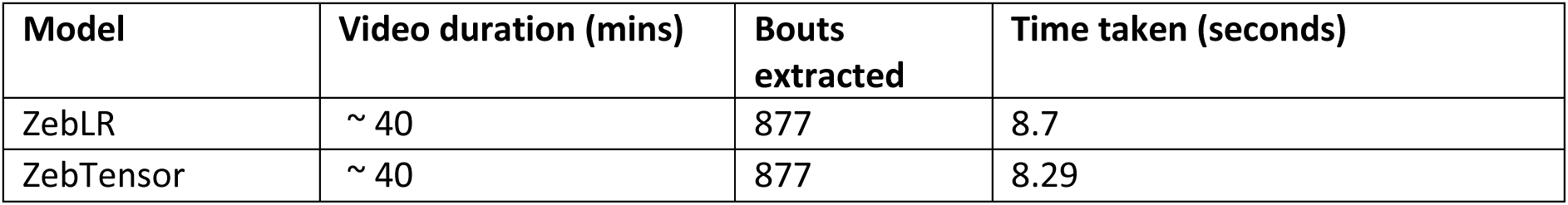
Example time taken for analysis of a 40-minute video containing 877 swim bouts extracted with PoseR. The time taken to extract swim bouts and classify them using PoseR models for long high-frame rate videos.

| Model | Video duration (mins) | Bouts extracted | Time taken (seconds) |
| --- | --- | --- | --- |
| ZebLR | ~ 40 | 877 | 8.7 |
| ZebTensor | ~ 40 | 877 | 8.29 |

PoseR is installable via pypi https://pypi.org/project/PoseR-napari/ and hosted on GitHub at https://github.com/pnm4sfix/PoseR. All data is available at the Zenodo repository at https://doi.org/10.5281/zenodo.7807968. As the field of action recognition advances, further improvements in neural network architecture will be implemented within PoseR. These strategies typically involve including more context of pose estimations, converting them to multidimensional heatmaps as input for three-dimensional convolutional neural networks, and including an RGB video layer (56). Viewing pose estimations as graph structures is a powerful approach and can be further extended by expanding the number of features associated with each body part node. For example, local video features at the location of each pose key point can be assigned to each node and included in the feature matrix for convolution. This would enable the network to learn the local video context in addition to spatial representation and estimation confidence of behaviors. Further, research into graph to text conversion and the use of language models to extract information from graphs is a very promising direction towards semantically describing behavior in an unsupervised way using only information contained in a pose graph or knowledge graph of a visual scene in video.

## Funding

This work was supported by the Biotechnology and Biological Sciences Research Council Grant BB/T006560/1, and an RS Macdonald Charitable Trust Grant. Manuscript revisions were supported by an Leverhulme Early Career Fellowship ECF_2022_105.

## Acknowledgements

We would like to acknowledge David Harris-Birtill for his invaluable advice, Cat Hobaiter and Stefan Pulver for their input and data-sharing during the conception and early stages of the project, and Jacqueline MacPherson, Michael Kinnear, James Lewis-Cheetham and Angus Aitken from the Psychology Workshop for their technical support. We’d also like to thank Joe Chapman and technical staff from the Scottish Ocean Institute for zebrafish husbandry support, and the University of St Andrews for computational support via its central HPC facility.

## Supplementary figures and figure legends

**Figure S1.**
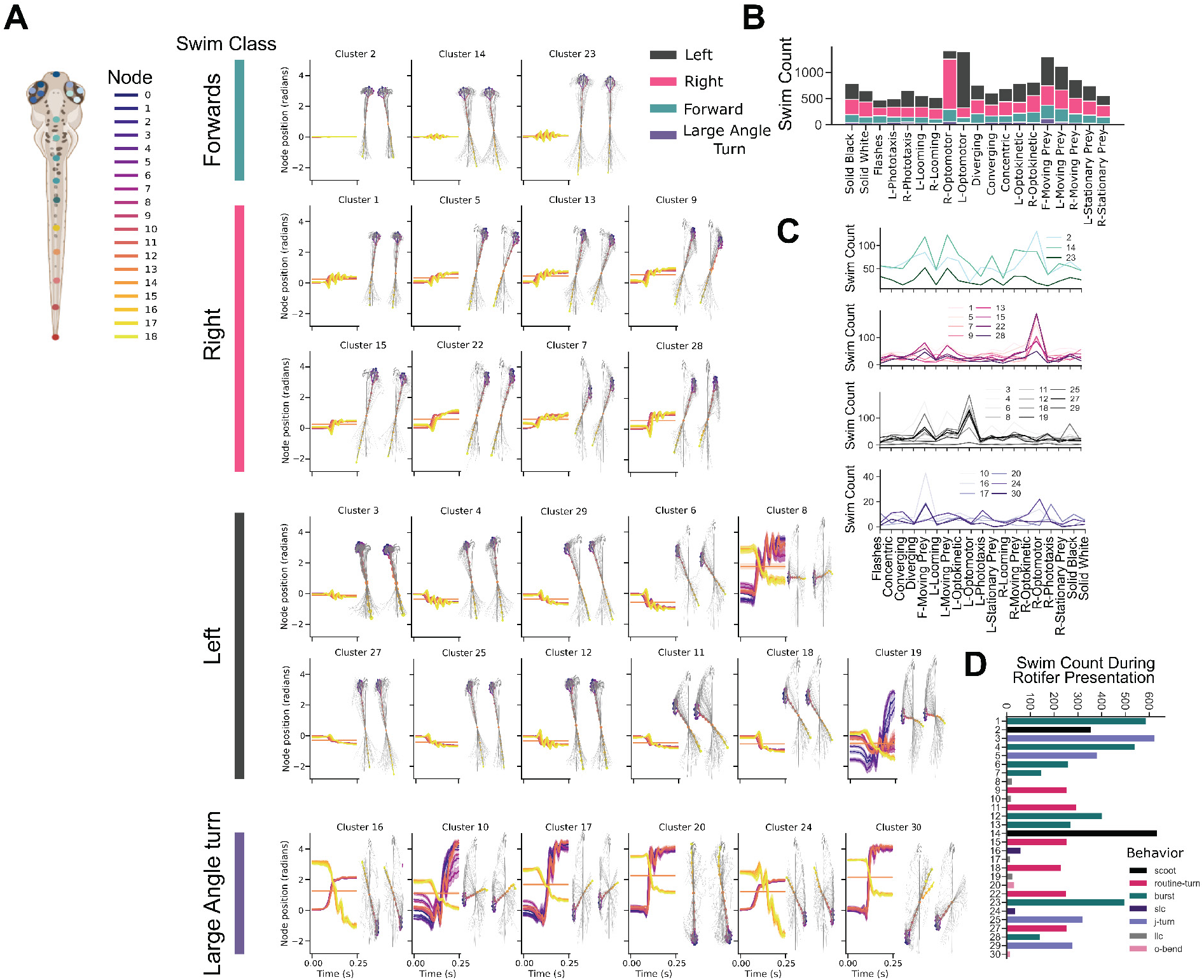
Swim dynamic and cluster examples. **A)** A schematic displaying the location of key point nodes along the zebrafish larval body. Each cluster (excluding 21 and 26 due to pose estimation noise) is represented by the average position of each color-coded node in radians during swimming. Error bands are shown as standard error of the mean position. B) A stacked bar chart quantifying numbers of each broad class (left, right, forwards and large amplitude swim type) according to the presentation of visual stimuli. C) Parallel plots showing the number of each swim type cluster across visual stimuli. D) Swim counts of each swim cluster during presence of live rotifers.

**Figure S2.**
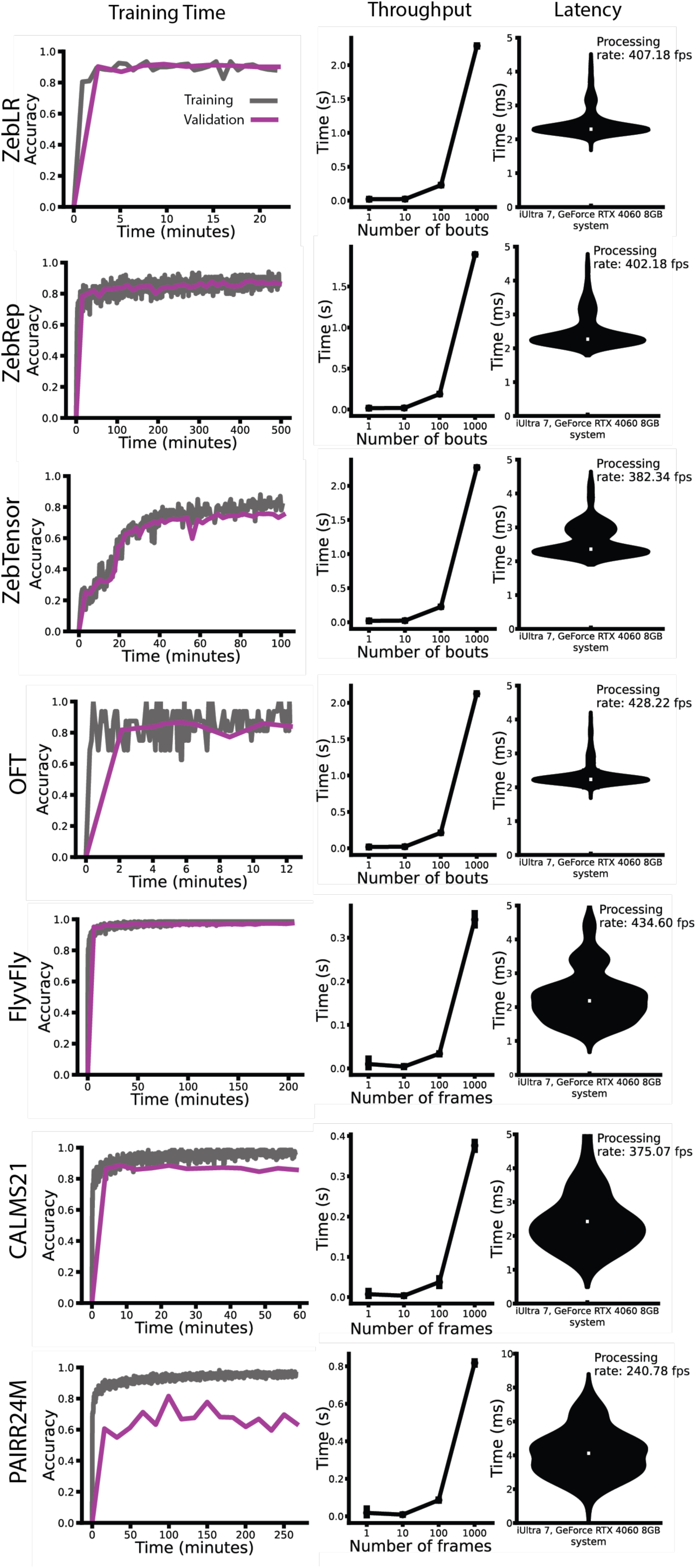
Training time, throughput and inference for all datasets. Left column: Example training time profiles representing training and validation accuracy as a function of time in minutes. Middle column: The time taken to process 1, 10, 100 and 1000 bouts or frames using a batch size of 10 for each dataset. Note that ZebLR, ZebTensor, ZebRep and OFT datasets were trained on extracted windows of behavioural bouts, whereas FlyvFly, CALMS21 and PAIRR24M were trained on frame-by-frame level annotations. Right column: The latency to process one frame or bout for all the datasets.

**Figure S3.**
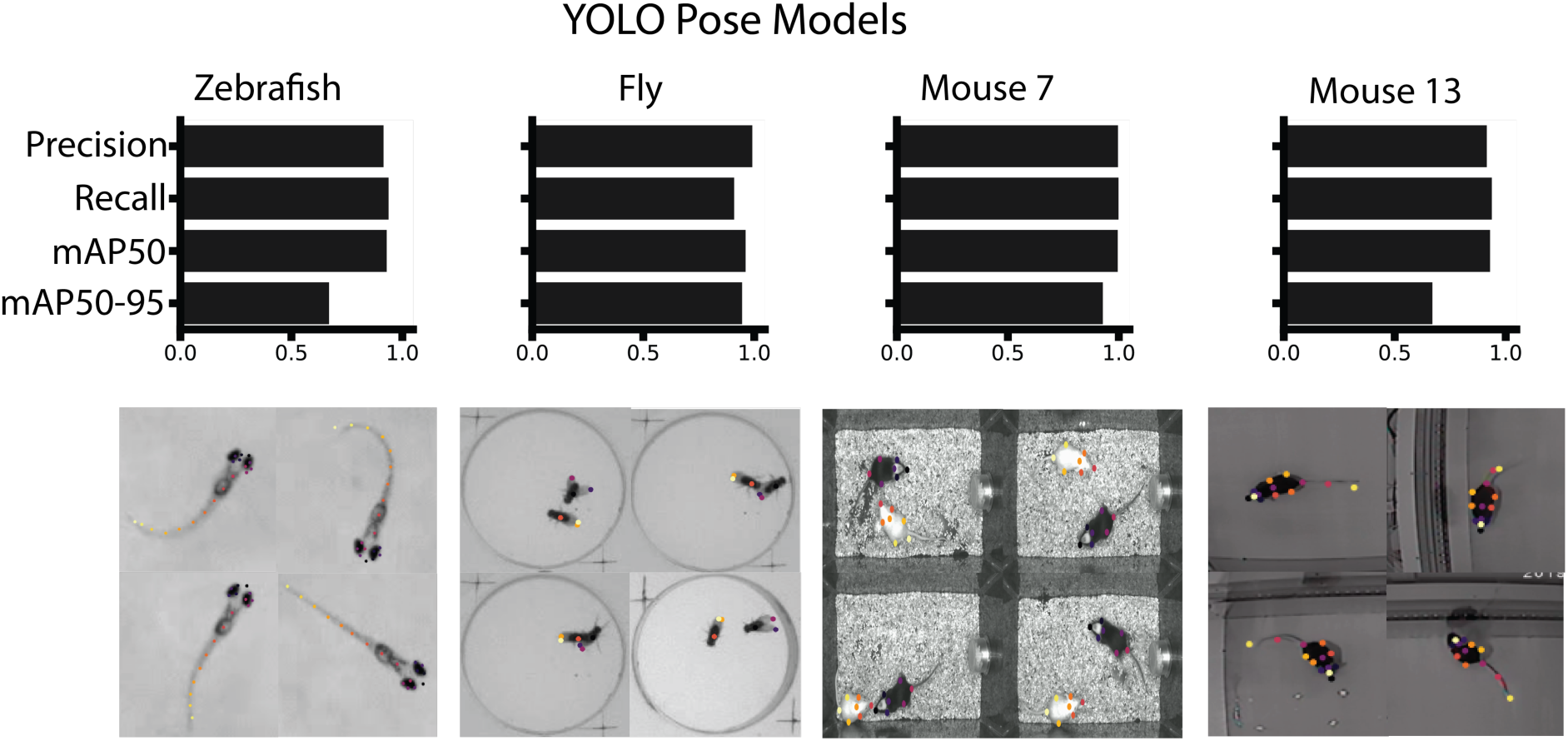
Pose estimations YOLO models and post-hoc correction in PoseR. Ultralytics YOLO pose estimation models available in the PoseR toolbox for each species. The zebrafish model pose estimations can be used as input to the zebrafish classifiers in PoseR, the fly model pose estimations can be used for PoseR classifiers trained on FlyvFly dataset. Mouse 7 pose estimations can be used for classifiers trained on CALMS21 social mouse dataset and Mouse 13 pose estimations can be used for classifiers trained on the open-field test dataset. Metrics reported are Precision, Recall, mAP50, mAP50-95 on the test set. mAP50 is the Mean Average Precision measured at a single IoU threshold of 0.5, assessing the model’s accuracy with at least 50% overlap, while mAP50-95 averages this precision over multiple IoU thresholds from 0.5 to 0.95, offering a more rigorous evaluation of detection accuracy across different overlap levels. Images reproduced for Fly, Mouse 7, Mouse 13 panels from FlyVFly dataset (35) published under CC0, CALMS21 dataset (37) and Sturman dataset (14) published under CC By 4.0.

## Notes

https://doi.org/10.5281/zenodo.7807968

### Competing Interest Statement

The authors have declared no competing interest.

### Summary of Updates

All text sections revised; figures updated; additional supporting information added in text and tables; additional analyses performed; additional behavioral contexts added.

